# Individual differences in spatial learning are correlated across cognitive tasks but not associated with stress response behaviour in the Trinidadian guppy

**DOI:** 10.1101/2020.05.19.103689

**Authors:** Pamela M. Prentice, Chloe Mnatzaganian, Thomas M. Houslay, Alex Thornton, Alastair J. Wilson

**Affiliations:** Centre for Ecology and Conservation, University of Exeter, Penryn Campus, Cornwall, TR10 9FE, UK

## Abstract

Cognition is vital for carrying out behaviours required for survival and reproduction. In animals, we now know that cognitive performance varies not just among species, but also among individuals within populations. While this variation is a prerequisite for contemporary natural selection, it is also true that selection does not act on traits in isolation. The extent to which cognitive traits covary with other aspects of phenotype (e.g. personality traits) is therefore expected to be an important factor in shaping evolutionary dynamics. Here we adopt a multivariate approach to test for spatial learning ability in a captive population of male Trinidadian guppies (*Poecilia reticulata*), and ask whether differences in cognitive performance are associated with (repeatable) differences in stress response behaviour. We focus on stress response for two reasons. First, functional links between cognitive traits and ‘stress coping style’ have been hypothesised. Second, individual-level studies of cognitive performance typically rely on multiple testing paradigms that may themselves be a stressor. Thus, there is a risk that variation in stress responsiveness is itself a cause of apparent, but artefactual variance in cognitive ability. Using a set of fish exposed repeatedly to two distinct spatial learning tasks (maze layouts), and an acute stress response test (open field trial), we find differences among-individuals in task performance that are repeatable within- and across maze layouts. On average performance improves with experience in the first maze, consistent with spatial learning, but not in the second. In both mazes, there is among-individual variation in the trajectory of mean performance with trial number suggesting individuals differ in ‘learning rate’. Acute stress response behaviour is repeatable but predicts neither average time to solve the maze nor learning rate. We thus find no support for among-individual correlation between acute stress response and cognitive performance. However, we highlight the possibility that cumulative, chronic stress effects may nonetheless cause observed declines in performance across repeats for some individuals (leading to lack of improvement in mean time to solve the second maze). If so, this may represent a pervasive but difficult challenge for our ability to robustly estimate learning rates in studies of animal cognition.

## Introduction

Cognition is defined as the set of mechanisms by which animals acquire, process, store and use information from the environment (Healy & Rowe, 2010; Shettleworth, 2010), and is vital for carrying out day-to-day behaviours needed for survival and reproduction. While differences in cognitive performance among-species have long been studied in comparative psychology (for a review see Healy 2019), a more recent focus in behavioural ecology has been the characterisation of among-individual variation within populations of non-human animals (Lucon-Xiccato & Bisazza, 2017a; Ashton et al., 2018; Boogert et al., 2018). This among-individual variation is interesting from an evolutionary perspective, as it is a pre-requisite for natural selection and genetic variation – both of which are fundamental for adaptive evolution to occur (Wilson et al., 2010). Though selection does not act on traits in isolation, so quantifying relationships between cognitive variation and other aspects of phenotype is important. For example, functional links between cognitive performance and other aspects of behaviour (including, for example neophobia, boldness and stress responsiveness) have been widely hypothesised (Quinn et al., 2012; Sweis et al., 2013; Griffin et al., 2015; Medina-García et al., 2017). However, robustly testing these relationships is often challenging, requiring multivariate data collection and analyses to detect and describe patterns of variation between associated traits at the appropriate level (e.g., among-individual and/or among genotype; Dingemanse & Dochtermann, 2013). Nonetheless, such efforts are important if we hope to understand the adaptive evolution of cognition in the context of the wider phenotype (Thornton & Wilson, 2015). Here we address this broad goal in the more specific context of testing hypothesised links between cognitive performance and a behavioural stress-response (Øverli et al., 2007; Gibelli et al., 2019) in Trinidadian guppies (*Poecilia reticulata*).

Quantifying patterns of among-individual variation in cognitive traits is still in its infancy (Rowe & Healy, 2014; Thornton et al., 2014; Boogert et al., 2018), and empirical studies therefore remain somewhat limited (but see Ashton et al. 2018; Tyrone Lucon-Xiccato & Bisazza 2017; Niemelä et al. 2013 for examples). However it is now abundantly clear that populations typically harbour high levels of among-individual variation in behavioural traits more generally (Dingemanse & Réale, 2005). Individual differences in (mean) behaviours, commonly referred to as personality, can manifest as, for instance, variation in aggressiveness or sociability towards conspecifics, or differences in response when faced with predators or other sources of perceived risk (Bridger et al. 2015; Réale et al. 2007). Since strong directional or stabilising selection is usually predicted to erode variation (Roff, 2002), it is hypothesised that variation in personality traits is maintained by fitness trade-offs of some kind (Dingemanse et al., 2004; Quinn et al., 2016). For example, bolder individuals may be better at acquiring resources to invest in life history traits (e.g., growth, reproduction) but their behaviour may also expose them to greater predation risk. In this way personalities can themselves be viewed as components of life history strategies, leading to an expectation that they will be correlated with – and trade-off against - other aspects of physiological, reproductive, and behavioural phenotype (Réale et al. 2010; Sih, Bell, & Johnson 2004; Wolf et al. 2007). Certainly, arguments that trade-offs can maintain variation in cognitive performance parallel explanations made for widespread presence of personality. These could be trade-offs among cognitive domains, or alternatively between – for instance – an overall cognitive performance trait (‘general intelligence’ (Plomin & Spinath, 2002; Galsworthy et al., 2005; Burkart et al., 2017)) and other aspects of the phenotype.

Variation in an animal’s stress physiology may provide one putative source of among-individual differences in both personality traits and cognitive performance (Raoult et al., 2017; Gibelli et al., 2019). The widely used concept of stress coping style model predicts that individuals will vary - both behaviourally and physiologically-along a proactive/ reactive continuum (Coppens et al., 2010; Koolhaas et al., 1999; Sih, Bell, & Johnson 2004). As originally posited, the model predicts proactive coping styles will express more ‘fight or flight’ type behavioural responses induced by adrenaline-response to stressors. At the other extreme, reactive coping styles will be more behaviourally ‘passive’ (e.g., freezing or hiding) and show high HPA(I) activity leading to cortisol response (Øverli et al., 2007; Carere et al., 2014). Various links to cognitive performance variation have been suggested. For instance, proactive styles are associated with ‘bold’, exploratory, risk-taking personalities that may present with more opportunities to learn initially. Conversely, greater behavioural flexibility associated with reactive coping styles (Coppens et al., 2010) may be important for tasks such as reversal learning, that require an ability to acquire (and use) new information under changing environmental conditions (Koolhaas et al., 1999; Sih & Del Giudice, 2012; Griffin et al., 2015). More generally, sensitivity to external stressors or challenges could impact performance in cognitive assays if more stressed individuals are simply more or less motivated and/or are focused on sources of risk rather than environmental cues of rewards.

Although hypothesised links between stress responsiveness (or coping style) and cognitive performance seem intuitive, empirical evidence is still limited to a small number of studies (Lukowiak et al., 2014; Mesquita et al., 2015; Bebus et al., 2016; Bensky et al., 2017; Brust & Guenther, 2017; Mazza et al., 2018). There are also contrasting studies in which either a weak or no relationship was detected (Cole et al., 2011; Carazo et al., 2014; Guillette et al., 2015). It is also possible that relationships are variable across different aspects of cognition. For instance in sailfin mollies (*Poecilia latipinna*), individual fish displaying less thigmotaxic behaviour (an anxiety related behaviour in fish) performed better in a discrimination learning task than highly anxious individuals, whereas the opposite was found in a reversal learning task (Gibelli et al., 2019). Clearly, there is need for more empirical work before a clear picture of the complex relationship between variation in cognitive performance and stress responsiveness/coping style is understood. Here we address this broad goal by testing the hypothesis that individual differences in cognitive performance and stress responsiveness are correlated in male Trinidadian guppies (*Poecilia reticulata*).

The guppy is a freshwater poeciliid fish that is widely used as a model in behavioural and evolutionary ecology. Methods for assaying among-individual ‘personality’ variation are well established in this species generally (Burns & Rodd, 2008; White et al., 2016), while guppies have been used in cognitive studies that target learning colour discriminations (Trompf & Brown, 2014; Buechel et al., 2018), numerical discriminations (Kotrschal et al., 2013; Lucon-Xiccato & Bisazza, 2017b), reversal learning (Buechel et al., 2018), spatial learning (Lucon-Xiccato & Bisazza, 2017c) and inhibitory control (e.g. Lucon-Xiccato & Bisazza, 2016). Here, we investigate the relationship between behavioural stress response and performance in a spatial learning task in which male guppies repeatedly navigated a maze to access females as a reward. The cognitive task was repeated using a second, differently structured maze in order that we could assess not just variation in learning within a single spatial context, but also ask whether – for instance – individuals displaying greater performance in trials using the first maze subsequently also performed better in the second. In the wild, male guppies usually utilize large home ranges during mate search and foraging (Croft et al., 2003), and as such spatial learning is expected to be an ecologically relevant trait (Brown et al., 2005). For our measure of stress responsiveness, we utilise ‘Open Field Trials’ (OFT). Widely used across species as a paradigm for characterising behavioural differences related to exploration, activity, and ‘shy-bold’ type variation (Gosling, 2001; Bell et al., 2009), previous studies on this captive population of guppies have highlighted the utility of OFTs for assaying behavioural stress response (see e.g., Prentice et al., 2020). Observed behaviours expressed in the OFT are both repeatable and plastic with respect to experimentally-manipulated stressor severity (specifically perceived predation risk) (Houslay et al., 2018). We also know from pedigree-based quantitative genetic studies that individual (mean) behaviours and their predictability (defined as within-individual variance) are heritable (White, 2019, 2019; Prentice, 2020). Furthermore, there is evidence of genetic integration between OFT behaviour and cortisol expression, strengthening the view that the OFT provides an appropriate assay of behavioural stress response (Houslay et al., 2019).

In what follows we: i) test for evidence of learning in naïve guppies repeatedly exposed to a spatial learning task (maze), ii) ask whether individuals differ in cognitive performance across repeated trials and if so; iii) whether performance in the first maze predicts performance in a second spatial context (i.e. reconfigured maze). We predict that time to complete the mazes (our proxy of cognitive performance) will, on average, improve with experience consistent with spatial “learning”, but that individuals will differ consistently in cognitive performance within each maze. We also predict that individual performance in the first maze will be positively correlated with performance in the second, suggesting stable differences in cognitive ability, although we acknowledge proactive interference (difficulty inhibiting memory or previously learnt associations; Shettleworth, 2009a) may affect performance in the second maze. Finally, iv) we test the hypothesis that individual differences in cognitive performance will be associated with differences in stress responsiveness. Although empirical evidence suggests potential relationships in both directions between stress responsiveness and cognitive performance, with the current absence of specific models, we make no *a priori* predictions about the sign of the relationship here.

## Methods

### Study site and housing

All behavioural assays were carried out on guppies from a captive population (derived from wild fish collected in the Aripo River, Trinidad in 2008) housed at the University of Exeter’s Penryn campus. Adult males (n = 64) were randomly sampled from the stock population, and housed in groups of 8 in separate home tanks (15 l, 18.5 × 37 × 22 cm) maintained at 23–24°C on a 12:12 light/dark cycle. The tanks shared a recirculating sump water supply which underwent a 25% water change once per week. All fish were fed to satiation twice daily on commercial flake food and live brine shrimp (*Artemia salina*) to control as much as possible for energetic and nutritional states prior to testing. We elected to focus on males only for several reasons. First, pilot studies showed a high occurrence of ‘freezing’ behaviour in females (relative to males) when introduced to the maze. While freezing can be a component of the behavioural stress response (Houslay et al., 2018), we considered that frequent occurrence during the cognitive assay would complicate data interpretation. Second, males show consistent sexual reproductive motivation towards females (Burns & Rodd, 2008), enabling the use of females as a ‘reward’ for males solving the maze (Kotrschal et al., 2015). Third, male guppies exhibit distinctive markings and colouration on body and fins. By recording and sketching these for each fish we were able identify individuals within groups without the need to subject individuals to invasive tagging.

### Ethics

This work was conducted under the auspices of the Animals (Scientific Procedures Act) out with approval of the University of Exeter research ethics committee, under licence from the Home Office (UK) (Licence Number PPL30/3256). Experimental procedures and behavioural assays were developed in accordance with the principles of the three Rs and ASAB guidelines (Buchanan et al., 2020) for use of animals. All periods of handling and emersion were kept to a minimum and only fish deemed healthy and exhibiting normal behaviour were used in trials. At the end of the experiment, fish were returned to a designated ‘retirement’ tank (containing females as well as males) and not used in any further experiments.

### Overview of behavioural testing scheme

We used a repeated measures approach to test for among-individual (co)variation in spatial learning performance and stress responsiveness. Spatial learning was first assessed by repeatedly trialling individuals in a maze apparatus (Figure 1). Each individual fish was tested once per day for 11 consecutive days with reduction in time to complete the maze interpreted as ‘learning’. This is consistent with previous studies using either time to complete an objective or to perform a particular task to investigate variation in cognitive performance among-individuals (Guillette et al., 2015; Lucon-Xiccato & Bisazza, 2016; Mazza et al., 2018; Zidar et al., 2018). We acknowledge that this interpretation strictly requires the implicit assumption that the contribution of any other factors to among-individual variation (e.g., motivation, energetic state, experience previous to the experiment; Rowe & Healy 2014) is negligible relative to differential cognitive performance. We attempted to mitigate against other sources of among-individual variation as far as possible using standardised housing and husbandry conditions.

**Figure 1.**
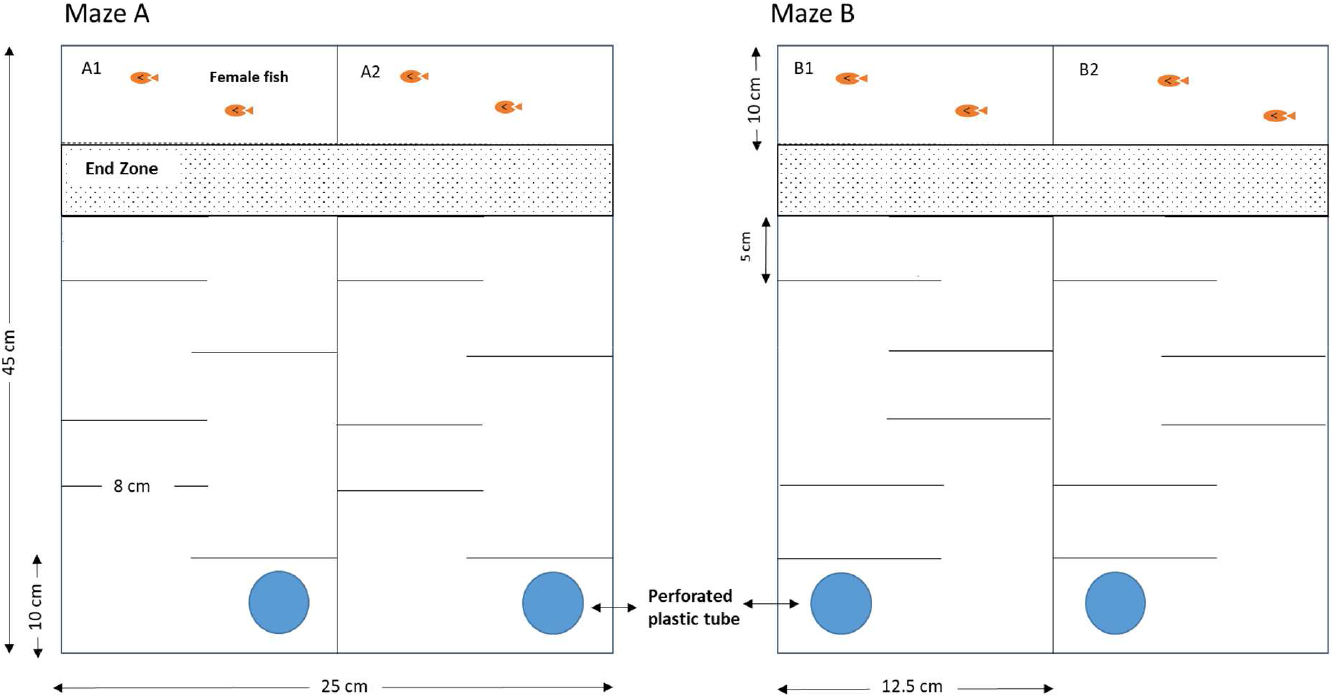
Aerial view of the maze designs used in the experiments (A and B), each tank was split into two identical mazes (1 and 2). Blue circles indicate perforated plastic tubes where individual fish started each trial, and time taken to reach the ‘end zone’ was recorded each trial.

Following completion of 11 spatial learning trials using Maze A, individuals were tested for stress responsiveness three times each over a three-week period using Open Field Trials (OFT) with a mean (range) of 4 (1-5) days between successive trials. Fish were then retested in a second maze (Maze B) with a different layout, with repeat trials conducted (as before) one per day for 11 consecutive days. Thus, in total, the design called for all individuals to complete 22 spatial learning trials, 11 on each of two different maze layouts (distributed across two different mazes) and three OFT over a total testing period of 43 days. Note that the sample size declined slightly across the experiment as (i) a few mortalities occurred naturally within the testing period and, (ii) we proactively ‘retired’ any fish not deemed to be feeding well and behaving normally in their home tanks as a precaution against cumulative adverse effects arising. Thus 63 fish experienced Maze A, which declined to n=60 at trial 11 and OFT testing. Five fish were then removed prior to experiencing Maze B (n= 55 at trial 1 and n=53 at trial 11).

### Spatial Learning Trials

In order to facilitate more rapid data collection, a single aquarium (25 x 45 x 25cm) was divided into two, with each half containing an identical version of maze A (A1, A2). Two replicates of maze B were similarly constructed (Figure 1). This allowed two fish to be tested concurrently during trials. Each maze consisted of 6 opaque Perspex panels (8 cm), spaced 5cm apart (Figure 1). A visually transparent perforated panel at one end of each maze was used to separate a small holding area (12.5 x 10 x 25 cm) contain two adult females selected randomly from stock. During trials the experimental maze tanks were lit from below by one fluorescent lamp and filled to a depth of 8 cm with room temperature water (approx. 23-24 °C). The water was taken from the same recirculating system used to house the male groups and was changed between each housing group (i.e. after every 4 runs with two fish trialled per run). Stimulus females were also changed at the same time.

At each trial, two males were individually netted from their home tank and quickly identified from natural markings. Each was randomly allocated to one of the two maze replicates and carefully placed within a perforated plastic tube at the start of the maze (Figure 1). They were given 60 s to acclimate before the plastic tubes were removed. A Sunkwang C160 video camera mounted above the tank allowed the fish to be observed without disturbance. Tracking software (http://www.biobserve.com) was used to determine *maze time*, measured as the total time taken to complete the maze after fish were released from the perforated plastic tube (with completion defined as reaching the ‘end’ zone; Figure 1). On reaching the ‘end’ zone, individuals were given 60 s undisturbed visual access to the females before an opaque plastic sheet was inserted to obstruct females from view. Following the 60 s reward period, fish were netted and returned to the home tank. To ensure standardized exposure to the reward stimulus, individuals that did not complete the maze within 480s of being released from the tube were gently guided through the maze to the end zone using a net behind them and then experienced 60 s visual access to the females. Following the 60 s reward period, fish were netted and returned to the home tank. These fish were assigned a right censored value of 480 second for *maze time.*

### Open Field Trial (OFT)

OFTs to characterise stress responsiveness closely followed the protocol described in White et al. (2016). For each trial, a single individual was netted from the home tank, quickly identified from natural markings and introduced gently into the centre of an open arena (a 30 × 20 cm tank filled to 5 cm water placed on a light-box). A cardboard screen around the tank prevented visual disturbance and a Sunkwang C160 video camera mounted above the arena again allowed movement to be tracked. Following a 30 s acclimation period, individuals’ movements were tracked for 4 minutes and 30 s to determine *track length* (total distance swum (cm)) and *area covered* (percent of tank area covered). These two observed behaviours which are known to be repeatable and heritable in this population (Houslay, 2018; White, 2019, 2019), were used to calculate the derived trait of *relative area* following Houslay et al. (2019). *Relative area* is the observed area covered in the trial minus the expected area covered under a simulated ‘random swim’ of length equal to the observed track length (see Houslay et al. (2019) for further detail on simulations). Low values of *relative area* result from a ‘flight type’ behavioural stress response in which individuals swim rapidly (yielding a high track length) but largely cover the same small area, thus covering relatively little area of tank arena. Low values of relative area are strongly correlated with thigmotaxis in this species, with fish swimming repeatedly across sections next to the tank walls and seeking escape from the tank arena. In contrast, high values of *relative area* correspond to efficient exploration (i.e. a high proportion of the arena covered given distance swum), by putatively less stressed fish.

### Statistical Analysis

Data from both types of behavioural assay were analysed using univariate and multivariate linear mixed effect models fitted by REML (restricted maximum likelihood) using ASReml within R (http://www.vsni.com) (Gilmour et al., 2009). By including individual identity as a random effect in these models we test for and characterise among-individual (co)variation. Traits were mean centred and scaled to standard deviation units to ease interpretation of results and facilitate convergence of multivariate models. For *maze time* we did this using the overall mean and standard deviation of observations from both mazes in order to preserve any differences in the distributions of performance times between A and B. With traits in standard deviation units (sdu), estimates of among-individual variance (V_ind_) can be interpreted as repeatabilities (i.e. proportion of the observed phenotypic variance explained by among-individual differences). However, we also calculate estimates of adjusted repeatability (R), the proportion of phenotypic variance explained by consistent among-individual differences, after controlling for fixed effects on the mean (Nakagawa & Schielzeth, 2010). Thus R=V_ind_/(V_ind_+VR) where V_R_ is the residual (within-individual) variance estimated from each model. The significance of random effects was tested using likelihood ratio tests (LRT), while fixed effects (included in the various models as described below) were tested using conditional F-statistics. All models assumed Gaussian error structures, deemed acceptable based on visual inspection of the model residuals.

### Univariate analyses of maze performance and spatial learning

We use *maze time* as our observed measure of performance. Here we describe in full the univariate analysis of data collected in maze A (subsequently *maze time_A_*). Identical procedures were then applied to data from maze B. First, we visualised the distribution of *maze time_A_* across repeat using box plots and also plotted the proportion of mazes completed as a function of repeat to see if a pattern of increasing average performance (i.e. decreasing *maze time* and/or increasing proportion of successful completion) was immediately apparent. Next a series of three nested models with identical fixed effects but differing random effect structure were fitted to the centred and scaled *maze time_A_* data. All models included a fixed effect of *trial number* (the cumulative number of trials experienced by an individual, treated as a continuous variable), allowing us to test for improvement in the mean (indicative of learning). Additional fixed effects were included as statistical controls for potential sources of variance not relevant to hypotheses being tested here. These included time of day (in minutes after 9 am), maze replicate (as a factor denoting position 1 or 2 in maze tank), and order caught from the home tank. The latter was to account for any cumulative disturbance effect of removing fish sequentially from the home tank and/or build-up of chemical cues in the maze between water changes.

The first univariate model of *maze time* contained no random effects, while the second contained a random intercept of individual identity. Likelihood ratio test (LRT) comparison of these models was conducted to test the hypothesis that individuals differ in their average performance (*maze time_A_*) across the 11 repeats, and we estimated the (adjusted) repeatability of performance under the second model. For the LRT we assume twice the difference in model log-likelihoods is distributed as a 50:50 mix of X^2^_1_ and X^2^_0_ following Stram & Lee (1994). The third model was a first order random regression (i.e. a random slope and intercept model) in which each individual’s deviation from the fixed effect mean *maze time* can change as a linear function of *trial number* (1-11). Variation in random slopes means that there is among-individual variation around the mean *maze time_A_* - *trial number* relationship. Thus, LRT comparison of the second and third models thus provides a test for among-individual variation in learning rate. This comparison is conducted assuming the test statistic is distributed as X^2^_2_ since the third model has two extra parameters (a slope variance and a slope-intercept covariance). Note that among-individual variance in slopes cannot be scaled to a repeatability as within individual variance in slope is not estimable (using data from a single maze; see below). Nor is its magnitude directly comparable to random intercept variance since slopes and intercepts are in different units. However, under the third model, among-individual variance in learning (slope) means that among-individual variance *maze time_A_* changes with *trial number* (Supplementary Information Figure S1). Thus, to understand the biological effect size of estimated variance in slopes, we use the third model to predict among-individual variance (V_ind_) and adjusted repeatability (R) of *maze time_A_* at both initial (trial 1) and final (trial 11) performance (following e.g., Nussey et al. (2007); see also Supplementary Information Table S3 for a didactic explanation of the linear algebra behind this). We note that among-individual variation at final performance has been used to infer differences in cognitive ability in studies adopting similar repeated measures designs (e.g. Langley et al. 2020) and so also has a useful biological interpretation here.

### Univariate analysis of relative area

To verify our expectation that individuals would show consistent differences in stress responsiveness, we fit a simple random intercepts model to (scaled and centred) *relative area.* This model included fixed effects of trial number (1-3), and time of day (in minutes after 9 am in which each trial took place) as well as a random effect of individual identity. Adjusted repeatability (R) of *relative area* was calculated and the significance of among individual variance tested by LRT comparison to a simplified model with no random effect (assuming the test statistic was distributed as a 50:50 mix of X^2^_1_ and X^2^_0_ as above).

### Multivariate modelling of Maze A, Maze B and OFT data combined

Finally, to test the predicted correlation structure between cognitive performance and stress responsiveness, we formulated a trivariate mixed model in which the three response variables were *maze time_A_, maze time_B_* and *relative area.* Fixed effects were exactly as described above on all three traits. Random effects were also as described above (i.e. individual level random intercepts and slopes for *maze time_A_* and *maze time_B_* but a random intercept only for *relative area*) but the multivariate formulation allowed us to estimate the full 5×5 among-individual covariance matrix (**ID**) among these effects. Since each observation of a fish provided data on a single trait only, residual covariances among traits were fixed to zero. After fitting the model, we compared it to a simplified fit in which all among-trait covariance elements in **ID** were constrained to zero. This provides a global test of individual covariance between traits. We then scaled estimated pairwise covariances in **ID** to their corresponding correlations for easier interpretation (noting for a pair of effects x,y the correlation rxy = COV_xy_/(V_x_V_y_)^0.5^. This allowed us to scrutinise the correlation structure between stress responsiveness and cognitive performance in both mazes A and B, using both final performance and learning rate (i.e. random regression slope) as measures of cognition.

This model also yields an estimate the individual level correlation in cognitive performance measures (final *maze time* performance, learning) across mazes. These are not strictly equivalent to individual repeatabilities of cognitive performance measures across mazes (as opposed to individual repeatability of *maze time* across trials within mazes) because estimates could be negative (Barbosa & Morrissey, 2021). However, they can be readily interpreted in those terms; a strong positive correlation between, for example, individual *learning* in maze A and maze B means this latent variable is highly repeatable across mazes. Conversely, a negative correlation means that individuals learning faster in maze A tend to learn more slowly in maze B (and *vice versa*).

## Results

### Performance in Maze A

Plots of the raw data suggest that average time to complete Maze A decreases across trials (Figure 2), and this pattern is qualitatively consistent with expectations if (average) performance improves as a consequence of learning. The mixed model analysis of *maze time_A_* confirms statistical support for this with a significant negative effect of trial repeat number (based on the full random slope and intercept model; coefficient = −0.058 (0.014) sdu, F_1,59.8_ = 17.890, P < 0.001. This effect size equates to an estimated decrease of 102.39 seconds in average *maze time* over the 11 trials. Other fixed effects of order caught and maze position were non-significant (see Supplementary Information Table S1). Likelihood ratio tests (LRT) confirmed among-individual variation in *maze time_A_* (comparison of null and random intercept models; χ^2^_0,1_ = 184.713 P < 0.001). Under the random intercept model, repeatability of *maze time_A_* conditional on fixed effects was estimated as R_A_ = 0.379 (0.052).

**Figure 2.**
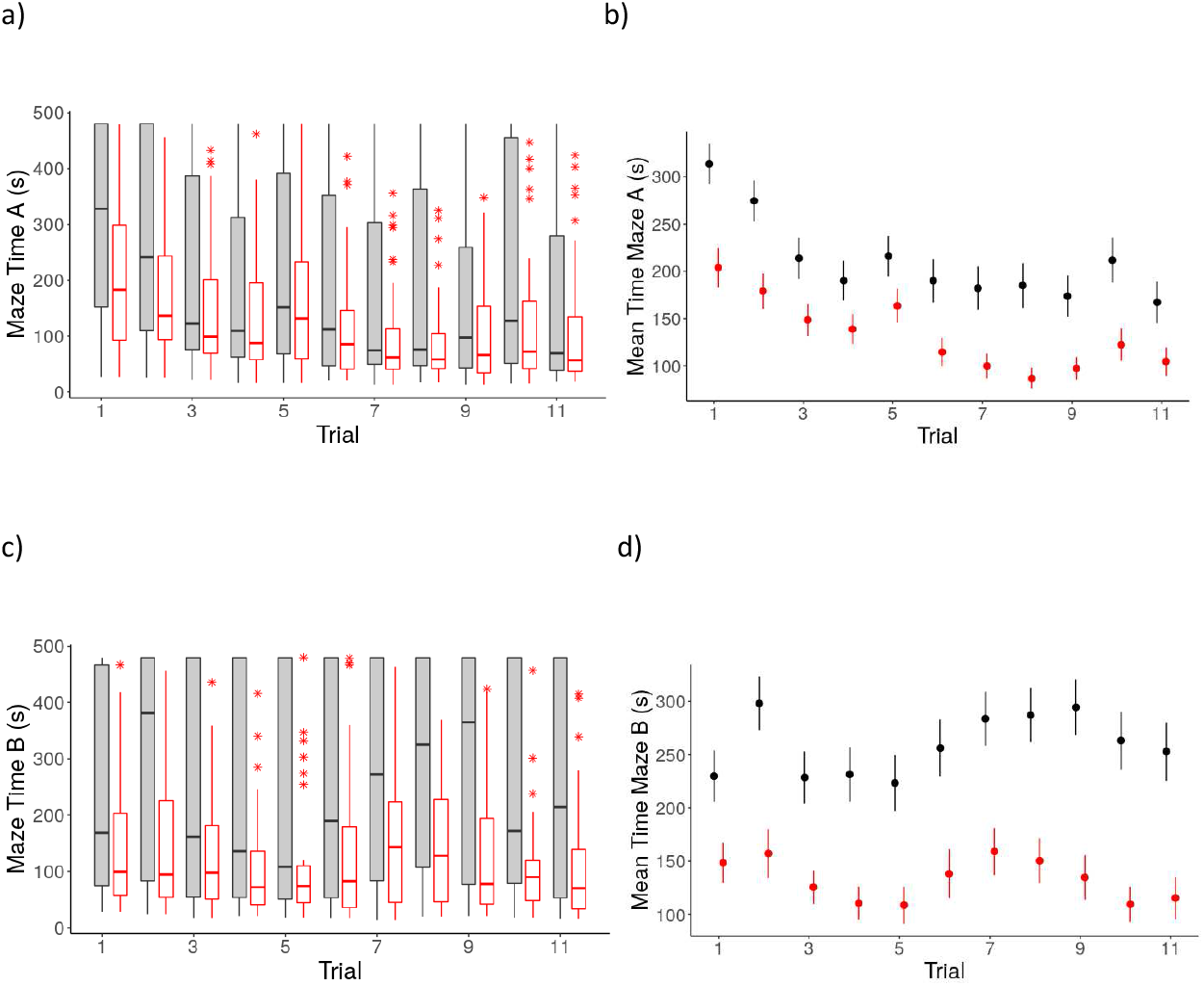
Plots of raw data of *maze time* across both maze designs. Boxplots (a) and (c) show the data distributions for time to complete Maze A and Maze B respectively across the 11 trials. Grey boxes display data of all individuals and red boxes represent only those individuals that successfully completed the task within 480 s. Horizontal lines within box correspond to behavioural medians, box boundaries correspond to first and third quartiles. When present, whiskers correspond to 10th and 90th percentiles, and points correspond to outliers. Plots (b) and (d) represent mean and standard errors for time to complete Maze A and B respectively. Colours represent the same groups; black error bars represent mean and standard errors of *maze time* for all individuals, and red represent only those individuals that successfully completed the maze in the allocated time.

LRT comparison of the random intercept and first order random regression models showed the latter to be a significantly better fit to the data (χ^2^_2_ = 26.990, P < 0.001). This comparison provides evidence for among-individual variance in the rate of change of *maze time_A_* across repeated trials. We interpret this (with caveats discussed below) as variation in the rate of spatial learning. Among-individual variance in intercepts (int) and slope (slp) were estimated as *V_ind_int__* = 0.402 (0.104) and *V_ind_slp__* = 0.006 (0.032) respectively while the among-individual intercept –slope correlation was estimated as (*r_ind_int_, ind_slp__* = −0.375 (0.168)). Biological interpretation of these parameters is not completely straightforward. Given the scaling of *trial number* in the random effect structure of the model (see Supplementary Information Table S3) *V_ind_int__* is interpretable as among individual variance in *maze time_A_* at first trial. While slope variance is in different units and thus not of directly comparable magnitude, variation in slopes actually means that among-individual variance in the observed trait (*V_ind_* for *maze time_A_*) changes with trial repeat number. Here the random regression model predicts values of *V*_*ind*_*A*1__ = 0.402 (0.104), and *V*_*ind*_*A*11__ = 0.677 (0.157) at first and last trial in maze A respectively, suggesting more among individual variation in performance at the end of trials than at the beginning. The negative intercept-slope correlation (*^r^_ind_A.int,A.slp__* = −0.375 (0.168)) means that individuals with higher intercepts (high *maze time_A_* at trial 1) tended to have lower (i.e., more negative) slopes indicative of faster learning. The corresponding predictions of repeatability at first and last observed trial are *R_A1_* = 0.429 (0.070) and *R_A11_* = 0.560 (0.063). These patterns are represented visually in Figure 3a, which shows the individual reaction norms predicted from the best linear unbiased predictions (BLUPs) of random intercept and slope for each fish (following e.g., Houslay & Wilson (2017)).

**Figure 3.**
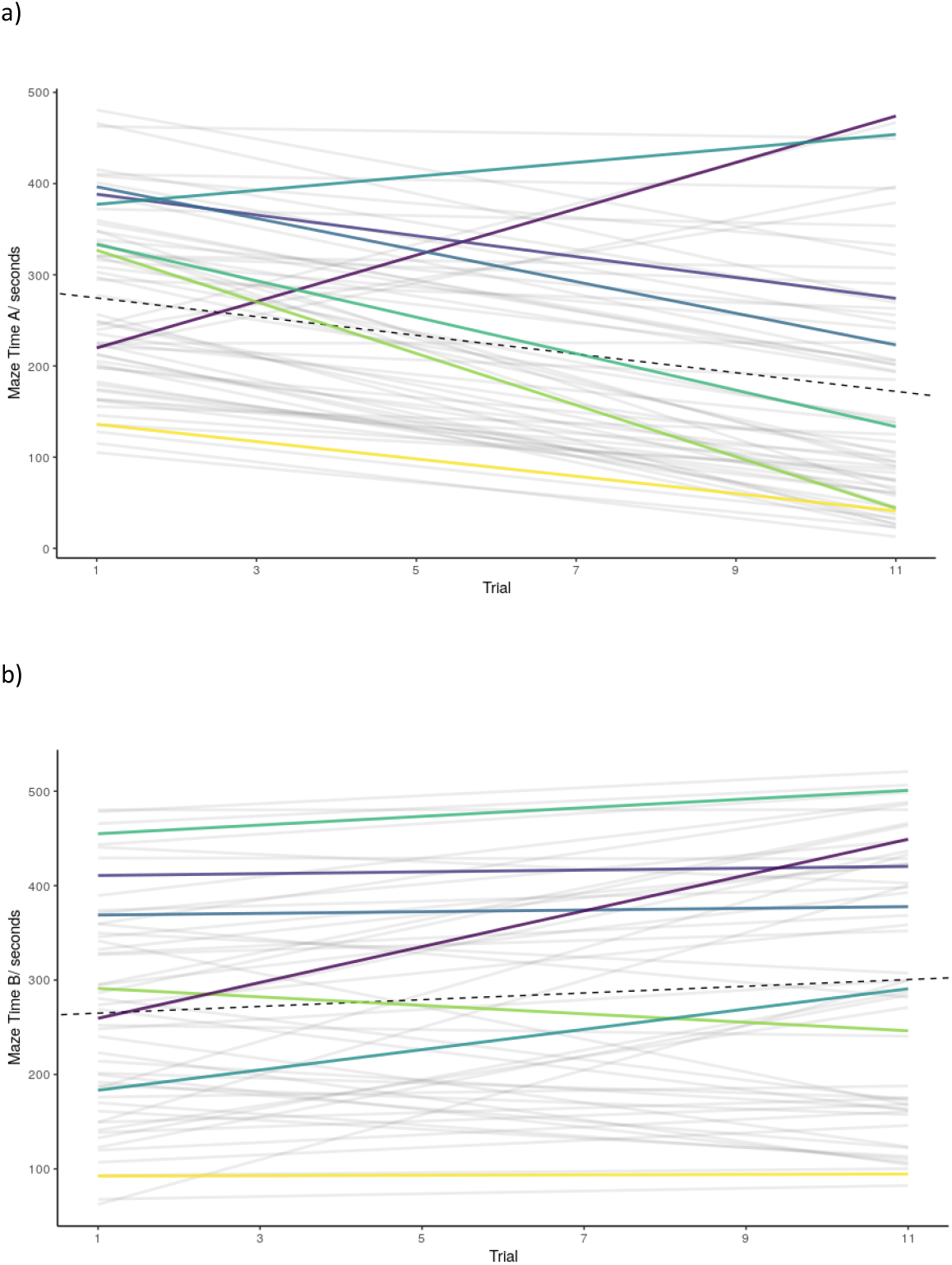
Spatial learning traits across Maze A and Maze B as a function of trial number, *maze time_A_* (a), and *maze time_B_* (b). Grey lines represent individual predicted reaction norms (BLUPs) from univariate random slope models for each trait. Coloured lines are used to illustrate reaction norms for a small random set of arbitrarily chosen individuals tested in both mazes (colours represent the same individuals across panels). The black dashed line represents the trend in fixed effect mean *maze time* across repeat trials

### Performance in Maze B

In contrast to Maze A, plotting *maze time_B_* data reveals no clear increase in performance (i.e. decrease in time) across trials. The mixed model analysis confirms the lack of improvement in the mean *maze time_B_*, with a (non-significant) positive estimate of the trial repeat number effect (from random slope and intercept model; coefficient =0.019 (0.014), F_1,53.8_ = 2.054, *P* = 0.175). Effects of order caught and maze position were not significant (Supplementary Information Table S1). Likelihood ratio tests (LRT) between the univariate random intercept model and the null model with no random effect, shows the presence of significant among-individual variation for *maze time_B_* (χ^2^_0,1_ = 200.048 P < 0.001), with a corresponding repeatability estimate of *R_B_* = 0.423 (0.056). The random slope model was a significantly better fit again (χ^2^_2_ = 15.926 P = 0.001) providing evidence of among-individual variation in the performance-trial number relationship. Among-individual variance in intercepts (int) and slope (slp) were estimated as 0.472 (0.125) and 0.005 (0.034) respectively. The intercept –slope correlation was negative as in Maze A (*r_ind_B.int,B.slp__* = −0.332 (0.190)). These estimates mean predicted values of *V*_*ind*_*B*1__ = 0.472 (0.125) and *V*_*ind*_*B*11__ = 0.659 (0.162)) which correspond to repeatabilities of *R_B1_* = 0.471 (0.072) and *R_B11_* = 0.554 (0.067). Although there is no (significant) effect of trial number on mean *maze time_B_* the presence of among-individual variance in slope suggest that some individuals are improving (consistent with learning) while for others performance is tending to get worse across repeats in Maze B (Figure 3b).

### Among-individual differences in OFT behaviour

We found evidence of significant among-individual variation in *relative area*, (*repeatability*(*with SE*), *R* = 0.465 (0.089), 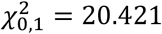, *P* < 0.001). This replicates previous findings in the same population (Prentice et al., 2020) though the current estimate of repeatability is somewhat higher, likely due to differences in study design (e.g. the current study used a shorter inter-observation interval and was limited to males only). Fixed effects from the OFT behaviour models are presented in the Supplementary Information Table S2 for completeness, although are not directly relevant to our hypotheses in this study.

### Multivariate model

The full multivariate model of *maze time_A_, maze time_B_* and *relative area* provides evidence of some significant covariance structure among traits at the individual level (comparison of the full model to one in which all among-individual between trait covariances are fixed to zero; *χ^2^_8_* = 48.844, P < 0.001). Examination of the estimated covariances and correlations (Table 1) suggests this result is largely driven by a strong positive correlation between the individual intercepts for *maze time_A_* and *maze time_B_* (*r_ind_A.int,B.int__* = 0.704 (0.127)). Biologically, this means that individual performance at first trial is strongly positively correlated across mazes. At final trial (i.e. 11), the individual correlation of performance across mazes is estimated at *r*_*ind*_*A*11,*B*11__ = 0.629 (0.119). Thus, our results show strong positive correlations of individual performance as measured by maze time across trials and mazes. This is not just true at first and last trial, but also for intermediate trial numbers within and across mazes (see Supplementary Table S3 for estimates and an explanation of their derivation from the random regression model output). However, taking a reaction norm interpretation of results, we do not find strong support for repeatable variation in learning rate across mazes. The individual correlation of reaction norm slopes is positive (*r_ind_A.slp,B.slp__* = 0.186 (0.246)), but relatively weak and not statistically significant (assuming approximate 95% confidence intervals of r±1.96SE). Nor do we find statistical support for among-individual correlation between maze performance intercepts or slopes (for either maze) and relative area (Table 1).

**Table 1.** Among individual variance–covariance–correlation matrix from the final trivariate model of *maze time_A_, relative area* and *maze time_B_*. Variances are shown on the diagonal (dark grey shading), with covariances above and correlations below. Light grey shading denotes within trait covariance/correlation estimates (i.e. between reaction norm intercepts and slopes). Standard errors are shown in parentheses and bold font denotes nominally significant pairwise estimates assuming approximate 95% CI of ± 1.96SE).

|  | *Maze time<sub>A</sub>* |  | *Relative area* | *Maze time<sub>B</sub>* |  |
| --- | --- | --- | --- | --- | --- |
|  | *intercept<sub>A</sub>* | *slope<sub>A</sub>* |  | *intercept<sub>B</sub>* | *slope<sub>B</sub>* |
| *intercept<sub>A</sub>* | **0.402 (0.104)** | -0.019 (0.012) | 0.096 (0.079) | **0.305 (0.089)** | -0.005 (0.010) |
| *slope<sub>A</sub>* | **-0.374 (0.168)** | **0.007 (0.002)** | -0.002 (0.011) | 0.005 (0.011) | 0.001 (0.001) |
| *Relative area* | 0.226 (0.178) | -0.034 (0.202) | **0.452 (0.118)** | 0.035 (0.086) | 0.012 (0.011) |
| *intercept<sub>B</sub>* | **0.704 (0.127)** | 0.094 (0.203) | 0.075 (0.186) | **0.467 (0.122)** | -0.016 (0.012) |
| *slope<sub>B</sub>* | -0.103 (0.226) | 0.186 (0.246) | 0.255 (0.221) | -0.335 (0.189) | **0.005 (0.002)** |

## Discussion

Here, we show evidence of among-individual differences in performance – measured as time to complete a maze – in guppies exposed to a spatial learning test paradigm. Performance of individuals is repeatable both within, and across, the two spatial learning tasks (i.e. mazes) presented. However, the question of whether there is robust evidence of learning, on average or by individual fish, is somewhat less clear cut. In particular, in the first maze used (A) we find evidence of improvement in mean performance consistent with learning (on average). We also find among-individual variation in this rate of improvement, and so – putatively – their rate of learning. However, the same fish exposed to maze B show (on average) no increase in performance across successive trials. We found among-individual correlation structure between performances (i.e. time in the maze) but not learning (i.e. rate of improvement) across the 2 spatial learning tasks. We did not find any significant association between individual differences in maze performance (or learning) and repeatable stress responsiveness as measured in the open field trials. In what follows we describe each of these findings in more detail and discuss them in the wider context of the cognitive literature.

The data from Maze A show that, on average, time to complete the maze improves across repeated trials. This improvement suggests that spatial learning is occurring in the guppies, a finding consistent with previous studies of this species (Kotrschal et al., 2015; Lucon-Xiccato & Bisazza, 2017c; Fong et al., 2019). We also see evidence of consistent, repeatable differences among-individuals in performance in Maze A. This is shown in our reaction norm models as significant among-individual variance in intercepts, which can be understood as performance at first trial. However, using among-individual variation in intercepts and slope to predict the corresponding variance at, and correlation among-, all trials (see Supplementary Information Table S3 for derivation and presentation of these estimates) reveals that in fact individual performance is positively correlated across all trials from 1 to 11. In simple terms, fish that are faster than average at completing Maze A in their first trial, tend to be faster than average across all subsequent trials too. Predicted repeatability of *maze time* is moderately high relative to many behavioural studies (e.g., 43% at trial 1, 51% at trial 11) but broadly comparable to estimates reported from similar assays designed to test cognitive variation (see Cauchoix et al., 2017 for an overview). We note that a contributing factor is likely to be short inter-observation period (here 24 hrs) typical of cognitive studies, since behavioural repeatabilities generally decline as this increases (Boulton et al., 2014).

Accepting that improvement across repeated trials can be interpreted as learning (caveats to this are discussed below), our random regression model also provides evidence for among-individual variation in spatial learning in Maze A. Usefully, our modelling strategy allowed all observations to contribute to estimating variance in the latent cognitive trait (learning) while avoiding statistically problematic ‘two-step’ analysis (Houslay & Wilson, 2017). Although this strategy is now widely used in studies of behavioural plasticity, it has not yet been widely adopted by researchers focussing specifically on animal cognition (but see e.g., Langley et al., 2020). In addition to finding variance in slopes (learning), we estimated a negative among-individual intercept-slope correlation using the Maze A data; individuals with higher intercepts (i.e. *maze time* at first trial) tend to have lower (more negative) slopes. While it is therefore the case that those fish performing poorly initially exhibit higher rates of learning, it is also true - as noted above - that individual performance (*maze time*) is positively correlated across trials 1-11. These two results are compatible because differences in learning (slope) are not sufficiently pronounced that initially poor performing (but fast learning) fish ‘overtake’ initially good performing (but slow learning) individuals by trial 11. We cannot comment on what fitness consequences, if any, the variation detected here would have in wild fish. Nonetheless, this finding does highlight a danger with any common presumptions that cognitive abilities may be under positive selection. Here, if we assumed that fitness benefits were accrued by rapidly achieving a spatial task (e.g. locating a resource) regardless of mechanism, it would be the slower learners that were advantaged. Thus, while it is tempting to assume fast learners will achieve better outcomes, they may sometimes simply be those with the ‘most room for improvement’.

Thus, findings from Maze A are consistent with our initial predictions that time to complete the maze would improve (on average) with experience due to spatial learning, but that individuals would also vary in both performance (*maze time*) and learning (rate of change in performance with experience). We also found that individuals that were quicker (over all trials) to complete Maze A, tended to be quicker (over all trials) to complete Maze B. While this could be attributable to cognitive differences, there are certainly other possibilities. For instance more explorative and/or less neophobic individuals may be generally faster at solving tasks (Boogert et al., 2006; Bousquet et al., 2015; Zidar et al., 2018). Similarly there could be among-individual variation in perceived cue salience (Meyer et al., 2012), individual physiology (Bókony et al., 2014), or motivation (van Horik & Madden, 2016). Regardless of these unknowns, an important difference between Maze A and Maze B was that we found no evidence of learning on average in the latter. In fact, for Maze B the mean *maze time* actually increased slightly, though not significantly, across trials. Thus, there is among-individual variation in intercept (*maze time* at trial 1) and also in slope. Given that there is no (significant) change in mean performance, but there is significant variation in slopes, we conclude that some individuals are improving (learning) in Maze B while others are getting worse with experience. We also note that, as in Maze A, slope variance is present, but not sufficiently high to break down the positive correlation structure of individual performance (*maze time*) across trials 1-11.

Although we did not formally test whether the average rate of learning across repeated trials differs between maze A and B it is reasonable to conclude it does (given no overlap of approximate 95% confidence intervals calculated as the slope ±1.96SE). Several possibilities may explain the finding of spatial learning on average in A but not B. First, the results from maze A may be a false positive (Sterne & Smith, 2001; Fraser et al., 2018). However coinciding with previous studies which show this species is capable of learning an initial spatial learning task (Kotrschal et al., 2015; Lucon-Xiccato & Bisazza, 2017b; Fong et al., 2019), we consider this unlikely. Second, it may be that the layout of maze B was more challenging to learn. This could certainly be true if, for instance learning to navigate a new maze following the acquisition of a previously learnt layout poses a more challenging task, for example due to proactive interference (difficulty inhibiting memory; Shettleworth, 2009a). In this case the second maze may require more trials to detect improvement. There is some evidence for such effects in guppies. For instance, Lucon-Xiccato & Bisazza (2014) found that on average guppies took 14.61 trials to learn a reversed colour cue association, while Fong et al., (2019) found that on average, 15.30 trials were required for guppies to learn a reversed maze layout.

A third possible explanation could be that learning is occurring in Maze B for some individuals, but that others experience some form or ‘trial fatigue’ or change in ‘state’ that reduces cognitive performance and/or motivation. For example, chronic stress effects may be arising from repeated capture and handling experienced in the experimental design (Huntingford et al., 2006; Warren & Callaghan, 1976; Wong et al., 2008). Such an effect could manifest in our Maze B data as an increase in the proportion of maze trials not successfully completed in <480 seconds (assuming trial fatigue increases the probability of not completing), coupled with a decline in average time across trial repeats for those trials that were successfully finished (assuming that better performing fish are learning). However, visual inspection of the data revealed no obvious trend in proportion of trials completed in <480 seconds, and post hoc analysis of maze B trials with censored records excluded provided no evidence of improvement either (results not shown). Empiricists obviously seek to minimise the possibility of chronic stress confounding conclusions from cognitive studies but it can be difficult to validate the implicit assumption that individuals remain (equally) ‘unstressed’ over experimental periods requiring repeated observations. Here we suggest this is a plausible hypothesis but not one we can currently test.

While the influence, or otherwise, of chronic stress on our results necessarily remains speculative, our experiment does confirm among-individual variation in acute stress behaviour as measured by *relative area* in the OFT. This replicates earlier results using independent data sets of fish from the same captive population (White, 2016; Houslay, 2019; Prentice, 2020). Acute stressor exposure can affect cognitive performance in spatial learning tasks in both mammals and fish (Gaikwad et al., 2011; Wong et al., 2019). At the individual level, there is also evidence that acute stress responses can predict outcomes under longer term chronic and/or repeated stressor exposure (Segerstrom & Miller, 2004; Salak-Johnson & McGlone, 2007; Øverli et al., 2007). However, we find no evidence of strong relationships between acute stress behaviour and performance or learning in either maze. Thus, we find no support for the prediction, made under the stress coping style model, that (acute) stress responsiveness will covary with cognitive performance (Coppens et al., 2010; Sih & Del Giudice, 2012; Griffin et al., 2015).

In summary, here we have evidence of consistent differences among-individuals in spatial task performance in the guppy *P. reticulata.* Individual performance is repeatable across trials within- and between two different spatial tasks (i.e. maze layouts). This among-individual variation in performance is consistent with underlying differences at cognitive factors but differences in ‘personality’ (e.g. neophobia, exploratory tendency) may also contribute. We also find evidence of improved performance with experience, consistent with spatial learning. In both tasks, variation around the trajectory of mean performance across trial number was present. While this means individuals can be considered as differing in ‘spatial learning rate’ it is important to note that performance declines for some individuals, especially in the second maze where there was no improvement in average time across 11 trials. We show here that an individual’s (repeatable) behavioural response to an acute stress stimulus does not predict either average performance in the maze or learning rate. However, we suggest the possibility that cumulative, chronic stress effects may contribute to declining performance (or reduced improvement) in our study. If individuals generally differ in susceptibility to chronic stress, this may represent a widespread but currently poorly acknowledged challenge for characterisation of cognitive variation in animal studies.

## Acknowledgements

We would like to thank technical staff in the Penryn Aquatic facility for help with technical and husbandry support. This work was supported by the Biotechnology and Biological Sciences Research Council (grant BB/L022656/1) and by a BBSRC studentship to PMP. We are also grateful to Neeltje Boogert and John Quinn for constructive comments on an earlier version of this MS.

## Notes

### Competing Interest Statement

The authors have declared no competing interest.

