## Supplemental Information for "Individual differences in spatial learning are correlated across cognitive tasks but not associated with stress response behaviour in the Trinidadian guppy"

### Supplementary Material

#### S1. Fixed effects from univariate models of *maze time*

Table S1: Fixed effect estimates from the full random intercept and random slope model for *maze time* in both maze A and B

#### S2. Fixed effects for univariate model of *relative area*

Table S2: Fixed effect estimates from the full random intercept and random slope model *for relative area*.

#### S3. Character state approach to visualising data

Table S3: Character state representation of among-individual correlation structure between trial specific *maze time* (both mazes) and *relative area*.

**Figure S1: Characterising individual variation in learning performance**

**Table S1**: Fixed effect estimates (with standard errors in parentheses) from the full random intercept and random slope model for *maze time* in both maze A and B

| Maze | Fixed effect | Effect size (SE) | DF | F | P |
|---|---|---|---|---|---|
| A | Intercept | 0.355 (0.119) | 1, 115.1 | 1.137 | 0.000 |
| | Trial | -0.058 (0.014) | 1, 59.8 | 17.890 | 0.000 |
| | Maze position (top) | 0.122 (0.062) | 1, 599.1 | 3.950 | 0.047 |
| | Order | -0.029 (0.015) | 1, 631.5 | 4.153 | 0.057 |
| B | Intercept | 0.031 (0.135) | 1, 102.8 | 0.001 | 0.563 |
| | Trial | 0.019 (0.014) | 1, 53.8 | 2.054 | 0.175 |
| | Maze position (top) | 0.071 (0.066) | 1, 535.4 | 1.163 | 0.281 |
| | Order | -0.378 (0.135) | 1, 559.7 | 4.949 | 0.034 |

**Table S2.** Fixed effect estimates (with standard errors in parentheses) from the full random intercept and random slope model for relative area from the open field trials (OFT).

| Model | Fixed effect | Effect size (SE) | DF | F | P |
|---|---|---|---|---|---|
| OFT | Intercept | 0.229 (0.451) | 1, 159 | 0.003 | 0.613 |
| | Trial | -0.262 (0.066) | 1, 116.8 | 15.710 | < 0.001 |
| | Time | 0.000 (0.000) | 1, 150.3 | 0.003 | 0.953 |

**Table S3.** Character state representation of among-individual correlation structure between trial specific *maze time* (both mazes) and *relative area*.

Table 1 in the main text presents the estimated among individual (**ID**) covariance matrix of reaction norm (RN) intercepts and slopes for *maze time*$_A$, *maze time*$_B$ and *relative area* (intercept only). Assuming the assumption of linear reaction norms hold true this can be transformed to the corresponding ‘character state’ (CS) among-individual covariance matrix of trial specific maze times and *relative area* (designated $\mathbf{ID}_{CS}$).

For a single trait (e.g. *maze time*$_A$), $\mathbf{ID}_{CS} = \mathbf{Q} \cdot \mathbf{ID}_{RN} \cdot \mathbf{Q}^T$ (following e.g. equation 5.8 in Roff et al., 2014), where $\mathbf{ID}_{RN}$ is the 2x2 covariance matrix of reaction norm (RN) intercepts and slopes, $\mathbf{Q}^T$ is the transpose of matrix **Q**, and **Q** itself contains the values of the covariate (trial number) at which we wish to evaluate $\mathbf{ID}_{CS}$. Where we want $\mathbf{ID}_{CS}$ to be an 11x11 matrix containing the among-individual variance in maze time at each trial number (1-11) on the diagonal, and the covariance between each pair of trial numbers in the off diagonal elements

$$\mathbf{Q} = \begin{bmatrix} 1 & 1 \\ 1 & 2 \\ 1 & 3 \\ 1 & 4 \\ 1 & 5 \\ 1 & 6 \\ 1 & 7 \\ 1 & 8 \\ 1 & 9 \\ 1 & 10 \\ 1 & 11 \end{bmatrix}$$

Following this, but expanded to the multivariate case, we transformed the estimated covariance matrix (**ID**) formulated under the trivariate model described in the main text (i.e., with individual effects on Maze A and B modelled as first order random regressions of trial number) to the corresponding character state matrix. This was then rescaled to yield point estimates of the among-individual correlation between trial specific performance within- and across-mazes, and between these performances and stress responsiveness. For simplicity we do not similarly attempt to transform estimates of uncertainty, but note that this table is a mathematical consequence (and transformation) of the statistical estimated presented in Table 1 of the main text (i.e. this is same set of results presented a different way).

| | A1 | A2 | A3 | A4 | A5 | A6 | A7 | A8 | A9 | A10 | A11 | RA | B1 | B2 | B3 | B4 | B5 | B6 | B7 | B8 | B9 | B10 | B11 |
|---|---|---|---|---|---|---|---|---|---|---|---|---|---|---|---|---|---|---|---|---|---|---|---|
| A1 | | | | | | | | | | | | | | | | | | | | | | | |
| A2 | 0.992 | | | | | | | | | | | | | | | | | | | | | | |
| A3 | 0.967 | 0.991 | | | | | | | | | | | | | | | | | | | | | |
| A4 | 0.923 | 0.964 | 0.991 | | | | | | | | | | | | | | | | | | | | |
| A5 | 0.861 | 0.918 | 0.962 | 0.991 | | | | | | | | | | | | | | | | | | | |
| A6 | 0.787 | 0.858 | 0.918 | 0.964 | 0.991 | | | | | | | | | | | | | | | | | | |
| A7 | 0.706 | 0.788 | 0.863 | 0.924 | 0.968 | 0.992 | | | | | | | | | | | | | | | | | |
| A8 | 0.623 | 0.715 | 0.802 | 0.876 | 0.934 | 0.973 | 0.994 | | | | | | | | | | | | | | | | |
| A9 | 0.543 | 0.643 | 0.739 | 0.824 | 0.894 | 0.945 | 0.978 | 0.995 | | | | | | | | | | | | | | | |
| A10 | 0.468 | 0.574 | 0.678 | 0.772 | 0.852 | 0.913 | 0.956 | 0.983 | 0.996 | | | | | | | | | | | | | | |
| A11 | 0.400 | 0.511 | 0.62 | 0.722 | 0.810 | 0.880 | 0.932 | 0.966 | 0.987 | 0.997 | | | | | | | | | | | | | |
| RA | 0.226 | 0.231 | 0.233 | 0.230 | 0.223 | 0.212 | 0.199 | 0.184 | 0.169 | 0.154 | 0.141 | | | | | | | | | | | | |
| B1 | 0.704 | 0.746 | 0.779 | 0.798 | 0.802 | 0.792 | 0.770 | 0.740 | 0.706 | 0.671 | 0.635 | 0.075 | | | | | | | | | | | |
| B2 | 0.715 | 0.761 | 0.797 | 0.819 | 0.826 | 0.818 | 0.798 | 0.770 | 0.736 | 0.701 | 0.666 | 0.105 | 0.995 | | | | | | | | | | |
| B3 | 0.718 | 0.767 | 0.806 | 0.832 | 0.842 | 0.837 | 0.819 | 0.792 | 0.760 | 0.726 | 0.691 | 0.136 | 0.978 | 0.994 | | | | | | | | | |
| B4 | 0.712 | 0.764 | 0.807 | 0.835 | 0.848 | 0.846 | 0.83 | 0.805 | 0.775 | 0.742 | 0.709 | 0.165 | 0.949 | 0.976 | 0.994 | | | | | | | | |
| B5 | 0.698 | 0.752 | 0.797 | 0.828 | 0.844 | 0.844 | 0.831 | 0.809 | 0.781 | 0.749 | 0.718 | 0.193 | 0.908 | 0.946 | 0.975 | 0.994 | | | | | | | |
| B6 | 0.676 | 0.731 | 0.778 | 0.811 | 0.830 | 0.833 | 0.823 | 0.803 | 0.777 | 0.748 | 0.718 | 0.218 | 0.857 | 0.905 | 0.946 | 0.976 | 0.994 | | | | | | |
| B7 | 0.647 | 0.703 | 0.751 | 0.786 | 0.807 | 0.812 | 0.805 | 0.788 | 0.764 | 0.737 | 0.710 | 0.239 | 0.799 | 0.856 | 0.907 | 0.948 | 0.977 | 0.995 | | | | | |
| B8 | 0.613 | 0.669 | 0.718 | 0.755 | 0.777 | 0.785 | 0.78 | 0.766 | 0.745 | 0.721 | 0.695 | 0.257 | 0.736 | 0.801 | 0.861 | 0.912 | 0.952 | 0.980 | 0.995 | | | | |

| | | | | | | | | | | | | | | | | | | | | | | |
|---|---|---|---|---|---|---|---|---|---|---|---|---|---|---|---|---|---|---|---|---|---|---|
| B9 | 0.577 | 0.633 | 0.681 | 0.719 | 0.743 | 0.753 | 0.751 | 0.739 | 0.721 | 0.699 | 0.676 | 0.271 | 0.672 | 0.744 | 0.811 | 0.871 | 0.92 | 0.957 | 0.982 | 0.996 | | |
| B10 | 0.540 | 0.595 | 0.643 | 0.682 | 0.707 | 0.719 | 0.719 | 0.709 | 0.694 | 0.674 | 0.653 | 0.282 | 0.608 | 0.686 | 0.76 | 0.828 | 0.885 | 0.930 | 0.963 | 0.985 | 0.997 | |
| B11 | 0.503 | 0.557 | 0.605 | 0.644 | 0.67 | 0.684 | 0.686 | 0.678 | 0.665 | 0.648 | 0.629 | 0.290 | 0.547 | 0.630 | 0.710 | 0.783 | 0.848 | 0.900 | 0.941 | 0.969 | 0.988 | 0.997 |

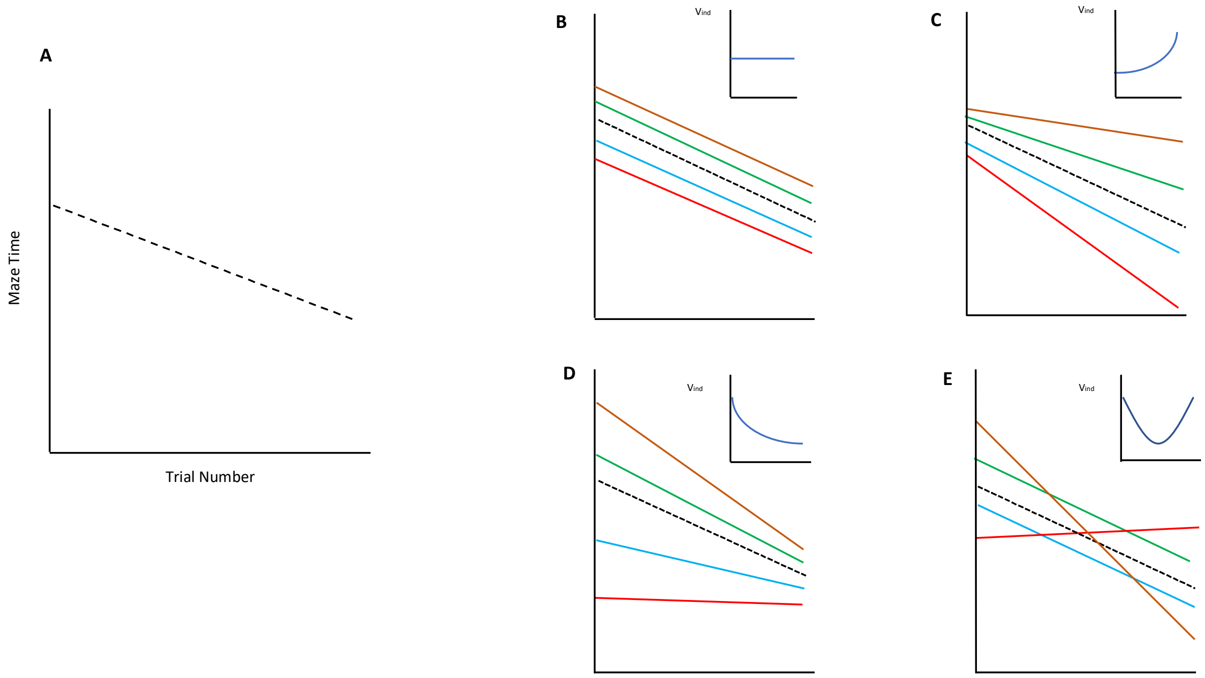


**Figure S1.** Characterising individual variation in learning performance. Main panel (A) shows an average (black dashed line) decrease in maze time with trial number from 1 to 11 consistent with learning. Inset panels show how individual trajectories may vary around this because of differences in reaction norm intercepts (B) and or slopes (C-E). Where slopes vary (C-E), a corollary of this is that the among-individual variance ($V_{ind}$) in maze time will change across trials. This could potentially increase (C) or decrease (D) monotonically, or there could be an intermediate trial number at which variance is minimised (E) or maximised (not shown). Where reaction norms tend to cross a lot within the range of trial numbers explored (E), this will result in low (and potentially negative) among-individual correlations between early and late trials.
